# Neglecting low season nest protection exacerbates female biased sea turtle hatchling production through the loss of male producing nests

**DOI:** 10.1101/752071

**Authors:** Catherine E. Hart, Luis Angel Tello-Sahagun, F. Alberto Abreu-Grobois, Alan A. Zavala-Norzagaray, Marc Girondot, Cesar P. Ley-Quiñonez

## Abstract

In the eastern Pacific, peak olive ridley sea turtle (*Lepidochelys olivacea*) nesting occurs during the warmest months which coincide with the rainy season, yet as nesting takes place year-round, the small proportion of the nests laid during dry-low season are exposed to contrasting environmental conditions. Most of the studies on Pacific coast sea turtles have estimated sex ratios produced during the rainy-high season when the majority of conservation activities take place. Thus, dry-low season nests have on the whole been overlooked. Here we compared sex ratios and hatchling fitness for offspring produced during the dry and rainy seasons during 2015. We found that protected olive ridley clutches incubated during the dry-low season were exposed to lower temperatures, yielded higher hatchling success, produced 100% male offspring and larger, heavier hatchlings with better locomotor abilities. Our results highlight the critical value of monitoring and protecting sea turtle nests beyond the peak season (when nests can be protected more efficiently) to include low season nests, albeit at much lower densities, but which by yielding higher proportions of males and with greater locomotor capacities may be the key to population viability and adaptation to anthropogenic climate change.

## 1. Introduction

Reproductive seasonality is present across species and phyla. Even in tropical regions where climatic variations may be less apparent, species maintain some level of seasonal pattern. In marine species, reproductive seasonality may be linked to marine productivity (Afán et al. 2015), local environmental features and large-scale environmental cues.

In the eastern Pacific, peak olive ridley sea turtle (*Lepidochelys olivacea*) nesting occurs during the warmest months which coincides with the rainy season. However, nesting can and does take place year-round, exposing the comparatively small number of nests laid in the dry and cold months to environmental conditions that contrast with those of the majority of nests incubating during the summer. For example, incubation temperature and humidity are markedly different between peak and low season. Temperature is one of the critical factors for the successful embryonic development of sea turtles (Miller 1985), in part because these species exhibit temperature-dependent sex determination (Mrosovsky and Pieau 1991; Broderick et al. 2000; Charruau and Hénaut 2012). Turtle embryos can develop a thermal tolerance range of between 25°C and 35°C (Howard, Bell, and Pike 2014). However, olive ridley clutches can survive higher temperatures (>37.9°C) but only over short durations but with detrimental effects on overall emergence success (Maulany, Booth, and Baxter 2012). Olive ridley turtles also present latitudinal variation in reported pivotal temperatures which produce 50 per cent of each sex within a clutch (Costa Rica: 31°C (Wibbels, Rostal, and Byles 1998); Mexico: 29.9°C (Sandoval, Gómez-Muñoz, and Porta-Gándara 2017). As incubation temperature rises above the pivotal within a sea turtle clutch, the proportion of females increases to a point of producing all females. The opposite is true as temperature falls below the pivotal and all-male production can occur in the lower viable temperature scale. Additionally, rainfall is a factor that varies greatly between seasons, especially in the tropics. Humidity within the nest environment influences moisture uptake by embryos, resulting in longer incubation durations and larger hatchlings (Delmas et al. 2007) and may also affect the sex ratio (Godfrey, Barreto, and Mrosovsky 1996; Wyneken and Lolavar 2015).

These factors make sea turtles particularly vulnerable to climate change (Fuller et al. 2013; Refsnider and Janzen 2016) which is predicted to not only cause increased incubation temperatures but also in sea level (IPCC 2007). Additionally, storms which are expected to become stronger and more frequent will further impact and modify turtle nesting habitat (L. Hawkes et al. 2009; L. A. Hawkes et al. 2013; Fuentes, Hamann, and Limpus 2010; Fuentes, Limpus, and Hamann 2011). Nonetheless, a female turtle can influence reproductive success through the choice of nesting location and depth at which she lays her eggs (David T. Booth and Freeman 2006; Santidrián Tomillo et al. 2017). However, even with the existence of female plasticity, sea turtles may have difficulty adapting to rapid climate change (L. Hawkes et al. 2009; Tilley et al. 2019). Olive ridleys may be the most adept of sea turtles to cope with environmental change due to their multiple reproductive strategies and observed flexibility in their degree of nesting site fidelity (Tripathy and Pandav 2007) and therefore may be able to choose sites that are less impacted by environmental change and which result in healthy offspring.

In recent years, phenotypical variation has been used to study the way changes in abiotic conditions affect hatchling fitness (Fisher, Godfrey, and Owens 2014; Liles et al. 2019). In warmer nests, hatchlings hatch sooner and consequently are smaller as less yolk is converted into tissue. Smaller hatchlings are slower during the crawl towards the ocean and during initial displacement from coastal zones (David T. Booth and Evans 2011) when compared with their larger counterparts. Larger hatchlings have the advantage of being too large a prey for certain predators. Furthermore, hatchlings must be able to maintain a 24-72 hour frenzied swimming period upon entering the ocean. Larger hatchlings which are stronger swimmers than smaller individuals could be more capable of avoiding the large aggregations of predators offshore of the nesting beach. Turtles in poor condition upon hatching have a reduced probability of avoiding predation (D. T. Booth et al. 2004; Wyneken and Salmon 1992; D. T. Booth 2009). The phenotype has also been used to evaluate the practice of nest relocation to hatcheries (Liles et al. 2019).

Since Mexico’s 1990 ban on sea turtle use and consumption, multiple nesting beach conservation programs have been created to protect clutches from illegal take and predation. However, due to limited resources, many sea turtle conservation projects are not able to continually monitor nesting beaches year-round. For species such as the olive ridley that nest along the Mexican Pacific, limits in resources forces conservation programs to focus on the rainy season months when nesting is significantly higher (García, Ceballos, and Adaya 2003), leaving nests laid during the dry season without protection. Dry season nests are often not monitored or counted leading to an impression from regional reports that nesting does not occur or is insignificant during this period. Registering dry season nesting is extremely important as their different abiotic conditions may affect hatchling sex ratio, phenotype, and fitness. Also, as sea turtle nesting seasons have been shown to shift in response to changes in temperatures (Weishampel et al. 2004; Pike et al. 2006; Witt et al. 2010), corresponding changes in phenology may be an adaptive strategy used by nesting turtles as a response to temperature increases due to climate change.

Nesting at Majahuas beach is part of the Playón de Mismaloya rookery which is notable for being the only known *arribada* rookery to collapse in the late 1970s due to high harvests of nesting females. The rookeries collapse resulted from a 99% reduction in nesting females (Abreu-Grobois and Plotkin 2008), which also resulted in the loss of genetic diversity (Rodríguez-Zárate, Rocha-Olivares, and Beheregaray 2013). Despite conservation efforts, there has been no *arribada* since the collapse. That said, solitary olive ridley nesting density is high in the area (García, Ceballos, and Adaya 2003).

Our goals were to 1) monitor nesting during a 12-month period; 2) compare incubation temperatures for nests incubated during the dry and rainy season; 3) determine if hatching success varied between these two seasons; 4) estimate sex ratios produced in monitored nests; 5) determine if incubation season had an effect on hatchling fitness and phenotypes; and, 6) discuss the conservation implications of the results.

## 2. Materials and methods

### 2.1 Study site

Majahuas beach is located in Jalisco between 19°50’41”N 105°22’40”W and 19°46’14”N 105°19’38”W on the Pacific coast of Mexico. Majahuas is southernmost 11 km of the Playón de Mismaloya sea turtle sanctuary. A RAMSAR mangrove wetland backs the beach.

The dry season lasts up to 8 months from November to June with a rainy season between July to October. Mean annual rainfall varies between 748 to 1000 mm with a mean temperature of 25°C (Bullock 1986).

### 2.2 Nest collection and incubation

We analyzed data collected by the fishing cooperative Roca Negra recorded during beach monitoring activities in 2015. Nests were protected via relocation to a hatchery (see below). We selected 71 nests at random (dry season: N = 37; rainy season: N = 34) for monitoring of incubation temperature and hatching success. Of these nests, 38 hatched and we conducted fitness tests on the hatchlings from these nests (dry season: N=28 nests; rainy season: N=10 nests).

Nests were collected during nightly beach patrols by either locating the recently laid nest via tracks or by encountering the nesting turtle. On encountering a female, we waited until she entered a trance-like state before taking morphometric measurements. Curved carapace length (CCL) and curved carapace width (CCW) were taken using a metric tape marked in 0.1 cm intervals. CCL was defined as the distance measured between the nuchal scute and the outer border of the post-central scutes and CCW was taken from the widest part of the carapace with the tape following the curvature of the carapace.

On locating a nest, the eggs were carefully removed from the egg chamber and counted. Nest depth was measured by placing a pole across the top of the mouth of the nest and the distance as taken from the pole to the bottom of the nest chamber. For each nest, we recorded the beach section and zone where it was laid (Intertidal (beach face to the berm) = A, Open beach (the berm to the vegetation line) = B and Beach (vegetation line to the dune) = C). Eggs were transferred to a plastic bag and transported to the hatchery using a quad bike located at km 2 of Majahuas beach. Each nest was reconstructed using a manual tree planter to achieve a standardized depth of 45cm and then the nest chamber was formed by hand to imitate the shape of a natural nest made by a female turtle. Eggs were transferred into the artificially dug chamber and a temperature logger (HOBO UA-001-08, Onset USA) was placed in the center of each clutch before being covered with sand. Temperature loggers measured 5.8 × 3.3 × 2.3 cm and were programed to measure the hourly temperature (accuracy of ± 0.5°C). Meteorological observations (daily maximum, minimum and mean air temperature) were obtained from the Universidad Autonoma de Mexico’s Biological Research Center in Cuixmala, Jalisco from 1 January 2015 to 31 December 2015.

### 2.3 Hatch lng phenotype and fltness

We selected 20 hatchlings at random upon emergence to partake in fitness tests and for phenotype measurements. When hatching success was too low to provide a total of 20 hatchlings, we conducted phenotype and fitness tests on those that were available. Hatchlings were weighed using an electronic balance (±0.1 g) and their straight carapace length (SCL), straight carapace width (SCW) and carapace depth were measured using an electronic calliper (±0.1 mm). Crawling speed (cm s^−1^) was recorded by measuring the time taken by each hatchling to crawl along a raceway of 3 m, 100 mm wide dug into the hatchery’s sand. We assigned hatchlings that failed to move within 300 s of being placed on the raceway to a failed to crawl category. We installed a LED light at one end of the raceway and care was taken to ensure that the track was flat. The time taken for hatchlings to self-right themselves was measured by placing the turtle upside down on their carapace and taking the time it took to right itself. This was repeated six times for each hatchling. If an individual took more than 60 s for any righting attempt, they were given a 5 s rest period on their plastron before the next attempt. After the tests, hatchlings were returned to the container with their siblings and then released into the ocean.

### 2.4 Sex ratio estimation

We used the R package ***embryogrowth*** v.6.4 (Girondot and Kaska 2014) to account for the effects of varying field temperatures on the dynamics of embryonic development and correctly identify the dates of the thermal sensitive period (TSP) when gonad differentiation occurred. With this, mean incubation temperatures during the true TSP were estimated for each nest and sex ratios derived using the thermal reaction norm for the species (Abreu-Grobois et al., in review). Sex ratio estimates are presented as mean ± SD unless stated otherwise.

### 2.4 Statistical analysis

Reported statistics are arithmetic means ± standard deviation (SD). All statistical were analyzed using Minitab® 18.1. (Minitab Inc., State College, Pennsylvania, USA). Kolmogorov-Sminov test was used as a normality test. Statistical test ANOVA with Tuckey’s method were carried out to examine mean differences among neonate fitness data obtained. A simple linear regression model for correlating the size of the adult females with number of eggs, and the incubation temperature with effect hatchling morphology. The *p values* of ≤0.05 were used to determine significant differences.

### 2.5 Ethics Statement

Permits were granted in Mexico by Dirección General de Vida Silvestre/Secretaría para el Medio Ambiente y los Recursos Naturales (SEMARNAT). Field Permits: SGPA/DGVS/05366/15.

## 3. Results

### 3.1 Nesting

We registered a total of 1954 nests over 12 months (1^st^ January 2015 - 30^th^ December 2015). Nesting occurred year-round with highest levels registered in October when the conservation project relocated 605 nests to the beach hatchery and lowest levels in May (n= 22 nests) (Fig. 1a). The majority of nests (n = 1573, 80.5%) were laid in rainy season while 19.5% of nests (n = 381) occurred during the dry season. Nesting was predominately on beach berm or zone B where 79.3% of nests were laid (n = 1547 nests) (zone A: 7.3%, 143 nests; zone C: 13.4%, 261 nests). Proportionally a greater number of nests were laid in intertidal zone C during the dry season (9.0% n=34 nests) than during the rainy season (6.9% n = 109). We measured 25 nesting females and found that the mean curved carapace length (CCL) and width (CCW) was 67.6 cm (range: 63–76 cm) and 73.8 cm (range: 68–82 cm), respectively (Supplementary table 1). There was no significant relationship between the size of the adult females and number of eggs laid (*R*^*2*^*=0.26; p=0.36*) or size of hatchlings.

**Figure 1.**
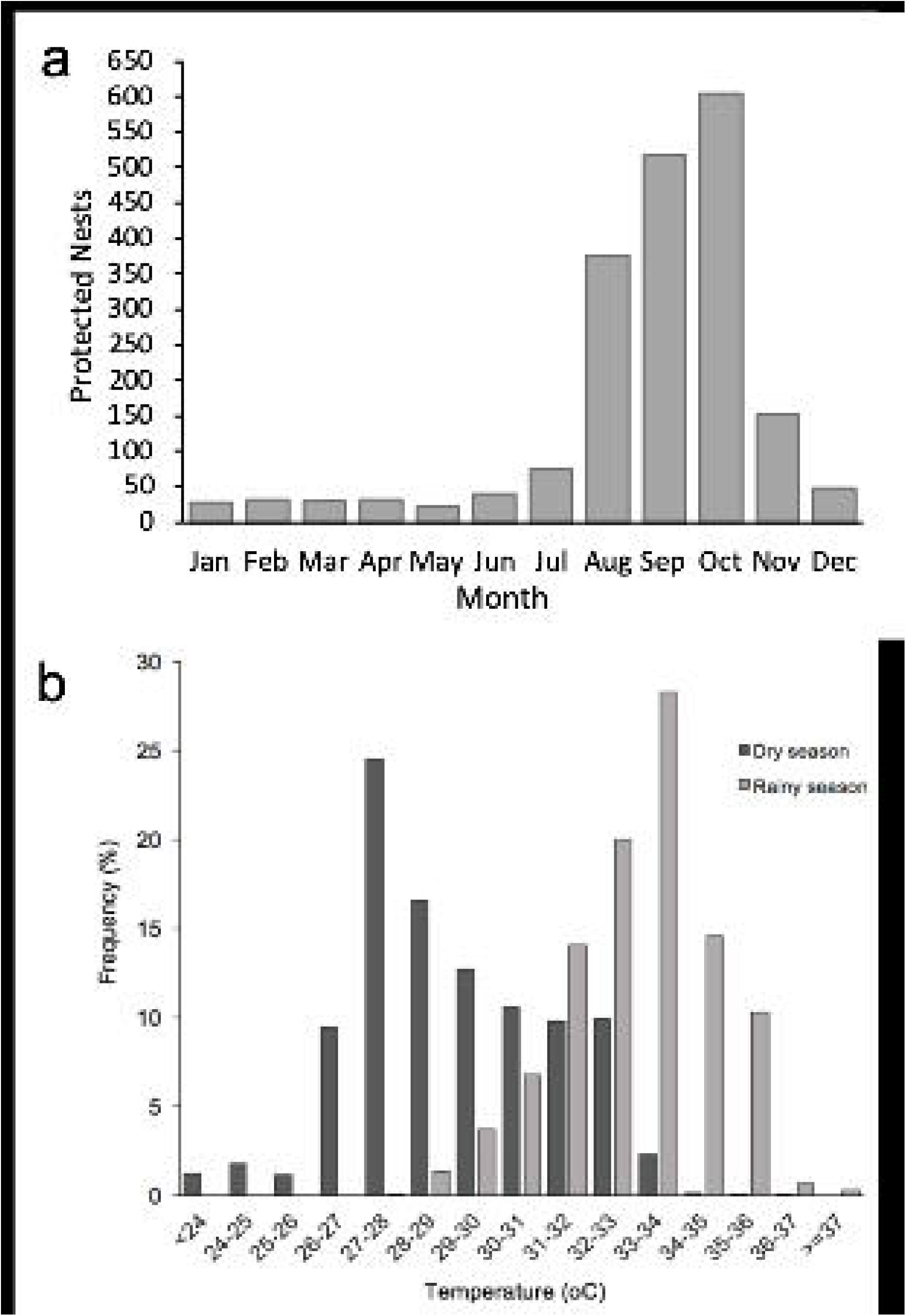
(a) Temporal distribution of nests protected at Majahuas beach during 2015 (b) Temperature frequency registered in the centre of hatched clutches during incubation.

### 3.2 Nest Temperature

Nest temperatures presented significant seasonal differences (*F*_*(1,69)*_*=143.26; p<0.001*) with those incubated during the dry season (29.09°C ± 0.52) being an mean of 3.89°C cooler than those incubated in the rainy season (32.98°C ± 0.58). Temperature within the 71 nests (Table 1) ranged between 22.8°C and 37.8°C. The most frequent temperature interval for dry season nests was 27-28°C with 24% of recorded values, while in the rainy season, the most frequent temperature interval was 33-34°C with 28% of records (fig. 1b).

**Table 1.** Results **s**ummary for data from 71 olive ridley clutches with ranges of incubation temperatures and estimated sex ratios (as proportion of males).

| Field code | Season | Starting incubation date | Incubation duration (d) | Clutch size | Hatching success | %>34°C | %<26°C | Mean °C ±SD (range) | Sex ratio |
| --- | --- | --- | --- | --- | --- | --- | --- | --- | --- |
| MJ1 | Dry | 18 Feb | 61.0 | 79 | 0.22 | 0.0 | 4.6 | 28.3 ± 1.6 (23.2-31.2) | 1.0 |
| MJ2 | Dry | 18 Feb | 60.1 | 87 | 0.95 | 0.0 | 4.6 | 28.9 ± 2.1 (23.6-32.9) | 1.0 |
| MJ3 | Dry | 18 Feb | 59.5 | 100 | 0.62 | 0.0 | 4.5 | 28.9 ± 2.1 (23.7-32.9) | 1.0 |
| MJ4 | Dry | 16 Feb | 62.0 | 104 | 1.00 | 0.0 | 4.5 | 28.8 ± 2.3 (23.2-33.2) | 1.0 |
| MJ5 | Dry | 16 Feb | 61.6 | 93 | 0.85 | 0.0 | 5.5 | 28.5 ± 2.3 (23.3-33.0) | 1.0 |
| MJ6 | Dry | 16 Feb | 60.4 | 103 | 1.00 | 2.6 | 4.4 | 29.2 ± 2.5 (23.8-34.7) | 1.0 |
| MJ7 | Dry | 16 Feb | 61.1 | 85 | 0.94 | 0.0 | 4.4 | 29.0 ± 2.2 (23.8-33.3) | 1.0 |
| MJ8 | Dry | 16 Feb | 60.3 | 96 | 0.85 | 0.0 | 4.6 | 28.8 ± 2.4 (22.8-33.3) | 1.0 |
| MJ9 | Dry | 16 Feb | 62.6 | 77 | 0.13 | 0.0 | 6.3 | 27.8 ± 1.6 (23.1-30.5) | 1.0 |
| MJ10 | Dry | 16 Feb | 59.9 | 92 | 0.92 | 0.0 | 4.7 | 28.9 ± 2.1 (24.0-32.8) | 1.0 |
| MJ11 | Dry | 19 Feb | 57.6 | 157 | 0.90 | 1.8 | 5.2 | 29.1 ± 2.7 (23.4-34.3) | 1.0 |
| MJ13 | Dry | 22 Feb | 62.5 | 87 | 0.67 | 0.0 | 6.1 | 28.5 ± 1.9 (24.2-31.5) | 1.0 |
| MJ14 | Dry | 24 Feb | 64.2 | 74 | 0.50 | 0.0 | 7.9 | 28.2 ± 1.9 (23.0-31.1) | 1.0 |
| MJ15 | Dry | 24 Feb | 59.1 | 123 | 0.89 | 0.0 | 6.2 | 28.8 ± 2.5 (24.0-33.1) | 1.0 |
| MJ16 | Dry | 25 Feb | 57.5 | 94 | 0.80 | 0.0 | 5.9 | 28.8 ± 2.3 (23.1-32.9) | 1.0 |
| MJ17 | Dry | 25 Feb | 57.5 | 96 | 0.85 | 0.0 | 5.1 | 29.1 ± 2.3 (23.9-33.2) | 1.0 |
| MJ18 | Dry | 25 Feb | 57.3 | 84 | 0.61 | 0.7 | 4.8 | 29.2 ± 2.5 (24.2-34.6) | 1.0 |
| MJ19 | Dry | 25 Feb | 60.8 | 113 | 0.85 | 0.0 | 4.5 | 28.8 ± 1.9 (24.4-32.0) | 1.0 |
| MJ20 | Dry | 25 Feb | 60.7 | 97 | 0.93 | 0.0 | 4.8 | 29.3 ± 2.4 (23.9-33.2) | 1.0 |
| MJ21 | Dry | 02 Mar | 58.1 | 102 |  | 0.0 | 6.3 | 29.0 ± 2.2 (23.6-32.4) | 1.0 |
| MJ22 | Dry | 02 Mar | 58.2 | 77 |  | 0.0 | 8.3 | 28.6 ± 2.2 (23.4-32.1) | 1.0 |
| MJ24 | Dry | 04 Mar | 63.5 | 92 | 0.90 | 0.0 | 7.5 | 28.8 ± 2.1 (23.5-32.0) | 1.0 |
| MJ25 | Dry | 04 Mar | 56.5 | 112 | 0.97 | 0.0 | 6.5 | 29.3 ± 2.3 (23.4-32.9) | 1.0 |
| MJ26 | Dry | 04 Mar | 56.5 | 79 | 0.81 | 0.0 | 5.3 | 29.3 ± 2.1 (24.3-32.2) | 1.0 |
| MJ27 | Dry | 04 Mar | 56.4 | 112 | 0.89 | 0.0 | 5.0 | 29.6 ± 2.3 (24.3-33.2) | 1.0 |
| **MJ28** | Dry | 04 Mar | 63.4 | 83 | 0.93 | 0.0 | 3.1 | 29.6 ± 1.9<br>(25.7-32.5) | 1.0 |
| **MJ30** | Dry | 05 Mar | 62.5 | 78 | 0.62 | 0.0 | 5.9 | 28.9 ± 1.7<br>(24.3-31.3) | 1.0 |
| **MJ31** | Dry | 19 Mar | 52.9 | 89 | 0.85 | 0.0 | 0.2 | 30.3 ± 1.6<br>(25.0-32.9) | 1.0 |
| **MJ32** | Dry | 21 Mar | 54.9 | 106 | 0.92 | 0.0 | 0.0 | 29.8 ± 1.7<br>(26.3-32.8) | 1.0 |
| **MJ33** | Dry | 21 Mar | 54.8 | 64 | 0.61 | 0.0 | 0.2 | 29.3 ± 1.3<br>(25.1-32.0) | 1.0 |
| **MJ34** | Dry | 21 Mar | 54.8 | 103 | 0.95 | 0.0 | 0.2 | 30.0 ± 2.0<br>(25.5-33.3) | 1.0 |
| **MJ35** | Dry | 21 Mar | 56.4 | 85 | 0.95 | 0.0 | 0.1 | 29.4 ± 1.8<br>(25.2-32.5) | 1.0 |
| **MJ36** | Dry | 21 Mar | 57.8 | 128 | 0.86 | 3.2 | 0.3 | 29.7 ± 2.3<br>(25.1-36.4) | 1.0 |
| **MJ37** | Dry | 21 Mar | 56.7 | 86 | 0.91 | 0.0 | 0.4 | 29.0 ± 1.9<br>(25.8-32.7) | 1.0 |
| **MJ38** | Dry | 22 Mar | 57.7 | 86 |  | 0.0 | 0.4 | 29.0 ± 1.4<br>(25.0-31.9) | 1.0 |
| **MJ39** | Dry | 22 Mar | 53.8 | 76 | 0.89 | 0.0 | 0.2 | 29.6 ± 1.7<br>(25.3-33.0) | 1.0 |
| **MJ40** | Dry | 22 Mar | 51.0 | 107 | 0.93 | 0.0 | 0.1 | 30.1 ± 1.9<br>(25.3-33.4) | 1.0 |
| **MJ42** | Rainy | 02 Jun | 47.7 | 92 | 0.54 | 41.1 | 0.0 | 33.5 ± 1.6<br>(29.5-36.5) | 0.0 |
| **MJ43** | Rainy | 02 Jun | 47.7 | 81 |  | 5.2 | 0.0 | 32.2 ± 1.1<br>(30.0-34.2) | 0.0 |
| **MJ44** | Rainy | 02 Jun | 47.7 | 98 |  | 0.0 | 0.0 | 32.0 ± 1.0<br>(30.2-33.6) | 0.0 |
| **MJ45** | Rainy | 03 Jun | 47.7 | 123 | 0.60 | 0.1 | 0.0 | 32.0 ± 1.2<br>(29.0-34.1) | 0.0 |
| **MJ47** | Rainy | 10 Jun | 50.8 | 121 | 0.92 | 8.2 | 0.0 | 32.3 ± 1.3<br>(29.7-35.2) | 0.0 |
| **MJ48** | Rainy | 20 Jun | 46.5 | 90 |  | 50.1 | 0.0 | 33.6 ± 1.5<br>(28.2-35.6) | 0.0 |
| **MJ49** | Rainy | 20 Jun | 46.7 | 109 | 0.93 | 1.5 | 0.0 | 31.8 ± 1.2<br>(28.3-34.1) | 1.0 |
| **MJ50** | Rainy | 21 Jun | 46.5 | 97 | 0.88 | 50.9 | 0.0 | 33.7 ± 1.5<br>(28.5-35.9) | 0.0 |
| **MJ51** | Rainy | 26 Jun | 42 | 80 |  | 3.1 | 0.0 | 31.9 ± 1.3<br>(28.4-34.3) | 1.0 |
| **MJ52** | Rainy | 26 Jun | 52.6 | 103 |  | 53.9 | 0.0 | 33.7 ± 1.6<br>(27.8-36.6) | 0.0 |
| **MJ53** | Rainy | 02 Jul | 46.1 | 93 | 0.69 | 40.0 | 0.0 | 33.7 ± 1.9<br>(28.8-37.8) | 0.0 |
| **MJ55** | Rainy | 02 Jul | 46.7 | 91 | 0.41 | 16.0 | 0.0 | 32.7 ± 1.5<br>(28.3-34.9) | 0.0 |
| **MJ57** | Rainy | 05 Jul | 47.6 | 98 |  | 40.4 | 0.0 | 33.4 ± 2.0<br>(28.1-37.7) | 0.0 |
| **MJ59** | Rainy | 05 Jul | 49.7 | 106 |  | 11.4 | 0.0 | 32.6 ± 1.5<br>(27.9-35.1) | 0.0 |
| **MJ63** | Rainy | 14 Jul | 46.2 | 105 |  | 17.1 | 0.0 | 33.5 ± 0.7<br>(30.4-34.9) | 0.0 |
| **MJ67** | Rainy | 14 Jul | 59.6 | 109 | 0.25 | 0.0 | 0.0 | 32.0 ± 1.4<br>(27.4-34.0) | 0.0 |
| **MJ69** | Rainy | 14 Jul | 59.6 | 111 |  | 2.8 | 0.0 | 32.5 ± 1.4<br>(27.9-34.4) | 0.0 |
| **MJ71** | Rainy | 23 Aug | 47.9 | 100 |  | 27.5 | 0.0 | 33.1 ± 1.5<br>(29.0-35.9) | 0.0 |
| **MJ73** | Rainy | 24 Aug | 47.8 | 87 |  | 28.8 | 0.0 | 33.2 ± 1.5<br>(29.0-36.0) | 0.0 |
| **MJ74** | Rainy | 24 Aug | 47.8 | 86 | 0.69 | 28.2 | 0.0 | 33.1 ± 1.6<br>(28.9-36.2) | 0.0 |
| **MJ75** | Rainy | 24 Aug | 44.8 | 56 | 0.88 | 0.4 | 0.0 | 33.9 ± 0.7<br>(31.2-36.1) | 0.0 |
| **MJ77** | Rainy | 24 Aug | Did not hatch | 68 | 0.00 | 0.3 | 0.0 | 33.2 ± 1.5<br>(29.0-36.7) | NA |
| **MJ78** | Rainy | 24 Aug | Did not hatch | 117 | 0.00 | 0.2 | 0.0 | 33.0 ± 1.5<br>(28.8-36.0) | NA |
| **MJ79** | Rainy | 24 Aug | Did not hatch | 120 | 0.00 | 0.3 | 0.0 | 33.2 ± 1.5<br>(29.2-36.0) | NA |
| **MJ98** | Rainy | 25 Sep | Did not hatch | 93 | 0.00 | 0.3 | 0.0 | 33.2 ± 1.8<br>(28.8-36.2) | NA |
| **MJ99** | Rainy | 25 Sep | Did not hatch | 66 | 0.00 | 0.4 | 0.0 | 33.3 ± 1.8<br>(28.6-36.3) | NA |
| **MJ100** | Rainy | 25 Sep | Did not hatch | 113 | 0.00 | 0.4 | 0.0 | 33.6 ± 2.0<br>(28.9-38.5) | NA |
| **MJ101** | Rainy | 25 Sep | Did not hatch | 87 | 0.00 | 0.3 | 0.0 | 33.1 ± 1.6<br>(29.1-35.9) | NA |
| **MJ103** | Rainy | 25 Sep | Did not hatch | 112 | 0.00 | 0.3 | 0.0 | 33.2 ± 1.5<br>(28.6-35.6) | NA |
| **MJ105** | Rainy | 25 Sep | Did not hatch | 92 | 0.00 | 0.2 | 0.0 | 32.8 ± 1.6<br>(28.3-35.3) | NA |
| **MJ106** | Rainy | 25 Sep | Did not hatch | 111 | 0.00 | 0.3 | 0.0 | 33.1 ± 1.7<br>(28.6-35.8) | NA |
| **MJ107** | Rainy | 25 Sep | Did not hatch | 98 | 0.00 | 0.3 | 0.0 | 33.2 ± 1.6<br>(28.9-35.8) | NA |
| **MJ108** | Rainy | 25 Sep | Did not hatch | 85 | 0.00 | 0.4 | 0.0 | 33.3 ± 1.7<br>(28.8-35.8) | NA |
| **MJ110** | Rainy | 25 Sep | Did not hatch | 93 | 0.00 | 0.4 | 0.0 | 33.5 ± 1.6<br>(29.5-36.0) | NA |

Within the hatchery, mid-nest depth temperature was lower than atmospheric temperature with tropical storms and hurricanes causing a visible drop in temperature (Supplemental fig. 1). However, mean incubation temperature within nests was not found to effect hatchling morphology (SCL (*R*^*2*^*=0.32964*), SCW (*R*^2^=0.05564), carapace depth (*R*^*2*^*=0.32831*) weight (*R*^*2*^*=0.06795*) or locomotor ability (righting propensity (*R*^*2*^*=0.01263*) righting time (*R*^*2*^*=0.01982*) or run speed (*R*^*2*^*=0.01415*)).

### 3.3 Sex ratio

We monitored the temperature inside 71 nests but were only able to estimate sex ratios in 57 due to 14 nests failing to hatch. All nests incubated in the dry season produced 100% male hatchlings, whereas those incubated during the rainy season (hatched nests: n=31) were female-biased with all but two nests producing 100% female offspring (Table 1).

### 3.4 Hatchling morphology and locomotor performance

Hatchling morphology was significantly affected by season, with dry season hatchlings presenting both larger SCL, (*Dry*: 40.62 mm ± 1.82; Rainy: 40.15 mm ± 2.53; *F*_*(1,758)*_*=7.16; p=0.008)* SCW (Dry: 32.84 mm ± 1.714; Rainy: 32.12 mm ± 2.104; *F*_*(1,758)*_*=20.71; p<0.001*) weight (Dry: 16.23g ± 1.686; Rainy: 14.93g ± 2.317; *F*_*(1,758)*_*=64.55; p<0.001*) than those hatched in rainy season. Significant differences in terrestrial locomotor performance were observed between seasons (*F*_*(1,758)*_*=60.17; p<0.001*), with dry season hatchlings having faster mean crawl speed (0.97 cm s^−1^ ± 0.594) than those hatched in the rainy season (0.55 cm s^−1^ ± 0.359). Rainy season hatchlings also presented slower mean righting response (3.33 s ± 2.11) than those hatched in the dry season (3.87 s ± 2.41) (Table 2). Overall hatching success was 52.7% and presented a significant difference between dry season hatchling success 74.3% and rainy season hatchling success 24.2% (*F*_*(1,1952)*_*=38.08; p<0.001*). Data for each nest studied can be found in Supplementary table 2.

**Table 2.** Mean temperature for 71 nests (37 in Dry season and 34 in Rainy season) and mean phenotype measurements (straight carapace length (SCL: mm), straight carapace width (SCW: mm) and weight (g)) and crawl speed and righting response for olive ridley sea turtle hatchlings from 38 nests at Majahuas beach by season (Rainy: n = 10; Dry: n = 28) in 2015.

| Parameter | Season |  |  |  | Statistical Test |
| --- | --- | --- | --- | --- | --- |
|  | Dry |  | Rainy |  |  |
|  | Mean ± SD | Min-Max | Mean ± SD | Min-Max |  |
| Temperature (°C) | 24.94 ± 1.858<br>(a) | 22.80-28.50 | 29.07 ± 0.807<br>(b) | 27.80-32.00 | $F_{(1,69)}=143.26$;<br>$p<0.001$ |
| Hatching Success (%) | 74.2 ± 2.97 (a) | 0.000-100.0 | 24.1 ± 3.56<br>(b) | 0.00-92.70 | $F_{(1,1952)}=38.08$;<br>$p<0.001$ |
| SCL (mm) | 40.62 ± 1.823<br>(a) | 34.00-47.50 | 40.15 ± 2.535<br>(b) | 30.00-49.00 | $F_{(1,758)}=7.16$; $p=0.008$ |
| SCW (mm) | 32.84 ± 1.714<br>(a) | 26.40-38.00 | 32.12 ± 2.104<br>(b) | 26.00-29.50 | $F_{(1,758)}=20.71$;<br>$p<0.001$ |
| Weight (g) | 16.23 ± 1.686<br>(a) | 12.00-24.00 | 14.93 ± 2.317<br>(b) | 8.020-19.88 | $F_{(1,758)}=64.55$;<br>$p<0.001$ |
| Crawl Speed (cm s<sup>-1</sup>) | 0.977 ± 0.594<br>(a) | 0.132-3.600 | 0.550 ± 0.359<br>(b) | 0.086-1.597 | $F_{(1,758)}=60.17$;<br>$p<0.001$ |
| Righting Response (s) | 3.870 ± 2.412<br>(a) | 0.980-19.00 | 3.336 ± 2.110<br>(b) | 0.830-18.00 | $F_{(1,758)}=04.641$;<br>$p=0.032$ |
| Righting Propensity | 4.471 ± 2.174<br>(a) | 0.000-6.000 | 4.598 ± 2.139<br>(a) | 0.000-6.000 | $F_{(1,758)}=0.352$;<br>$p=0.553$ |

N.B.: The statistical test used is the analysis of variance (ANOVA); statistical test data as mean ± SD followed by Tukey's test in parentheses if significant differences were found. Hatching success data in percentage.

## 4. Discussion

### Hatchling fitness

Seasonal effects were present in our study with dry season hatchlings having superior locomotor abilities and larger body size and weight than their rainy season counterparts. This is similar to other studies which have looked at the effect of nest temperature finding that cooler nests produce larger hatchlings (Booth, Freeney & Shibata 2013; Maulny et al 2013; Wood et al 2014) that may be better equipped (larger carapaces and flippers) to crawl and swim faster than their smaller counterparts from warmer nests (Ischer et al 2009). The phenotype and fitness advantages received from cooler incubation temperatures highlights the importance of protecting dry season nests which occur when nesting levels are low, as these nests produce have higher hatching success and the resulting hatchlings may have increased chance of survival as they may be quicker to exit predator rich coastal waters due to their larger size and better fitness characteristics.

Temperature is not the only factor that presents seasonal changes. Hatchlings entering the sea at different times of the year can encounter seasonal changes in oceanic circulation. Ocean currents can change in both intensity and direction. Therefore, neonates hatching at different times can end up in vastly different locations and be exposed to different conditions (Mansfield et al. 2017).

### Hurricane season

Hurricane season runs from May 15th to November 30th in the Eastern North Pacific (NHC), coinciding with peak olive ridley sea turtle nesting activity. The 2015 storm season was particularly active with 22 storms registered for the east Pacific of which 13 were hurricanes, six were tropical storms and three were tropical depressions. Seven storms (Supplementary Fig 1) affected Majahuas nesting beach during this study. This resulted in the loss of hundreds of nests due to beach erosion and wash-out of hatcheries. However, these storms also have the effect of lowering incubation temperatures, which help lower sand temperature in some cases below pivotal temperature. During August and much of September sand temperature remained above 34°C which has been identified as the lethal superior incubation temperature for some olive ridley populations. For example, when the effects of Hurricane Kevin and Linda occurred within the same week a fall in mid-nest depth temperature of 3°C (35°C to 32°C) occurred taking incubation temperatures out of lethal limits.

### Sex ratio

Dry season nests were estimated to be produce entirely male hatchlings, the increased hatch rate and survival of males may help balance out female biased sex ratios at Majahuas beach. Sandoval Espinoza (2012) estimated sex ratios for olive ridleys along the Mexican Pacific coast and found that ratios varied greatly with beaches in Jalisco (Chalacatepec and Playon de Mismaloya) producing 23% male sex ratios. For the Mexican Pacific, they estimated that temperatures would have resulted in male hatchling throughout the study period (July-Dec 2010) with 31% of males in September, 11% in August, 17% in October, 20% in November and 19% in December. They did not monitor temperatures in dry season. This is contrary to our results where the 2015 high rainy season temperatures resulted in very low levels of male hatchling production.

When we compare our results with those of a study in 1993 (Valadez González, Silva Bátiz, and Hernández Vázquez 2000) at a beach 5km north of Majahuas we find similar variations in sex ratio with 100% females produced in October and 100% males in December. However, the overall sex ratio of 7:3 in the 1993 in study is not the same as that found in Majahuas during our research. Incubation period in 1993 (Valadez González, Silva Bátiz, and Hernández Vázquez 2000) was 44 to 65 days, which is similar to our results where we recorded the longest incubation duration in February (64.2d) and the shortest in August (44.8d). However, temperature registered in the La Gloria beach hatchery ranged from 27°C ±0.10 (December) to 34°C ±0.36 (August) even when considering a higher temperature within nests due to metabolic heat (Sandoval et al. 2011) the 1993 study nests would not have experienced the extreme superior temperatures (max 37.8°C) that we registered within clutches. As expected from our 12-month study period, we registered lower temperatures than in the La Gloria study which did not monitor temperature during the dry winter season. Although Valadez-González et al. (2000) only recorded the hatchery sand temperature at 12-hour intervals at nest depth but not from within clutches, the study allows us to compare our results with data taken two decades ago.

### Benefits of low season nesting for females

Olive ridleys have been found to present the highest levels of multiple paternity in clutches than in any other sea turtle species, this is especially prominent in *arribada* breeding populations with 92% of nests having two or more fathers which led to the hypothesis that population size has a dominant effect on multiple paternity (Jensen et al. 2006). Yet the benefits of polyandry to female sea turtles have not been identified and multiple paternity was found to result in smaller clutches in Green turtles (Wright et al. 2013). Females that nest in times of low abundance are likely to encounter fewer males and therefore, benefit by a lower chance of multiple encounters with aggressive males (Jensen et al. 2006). However, small solitary breeding populations have also been found to present high levels of multiple paternity (Duran et al. 2015) and this could be a result of the low breeding and feeding site fidelity (Plotkin 2010) as well as a result of sea turtle females’ ability to store sperm for over a multiple years. During surveys in the Mexican Pacific in 2010, (Zepeda-Borja et al. 2017) observed olive ridleys mating only during October (autumn).

### Implications of current conservation effort

Concentrating effort and money on the peak nesting season may seem the best use of limited funds however nests laid during peak nesting season have lower possibilities of hatching than those laid in the low season due to lethally high temperatures and beach erosion due to storms. When protected from predation in hatcheries, the comparatively small number of nests laid in low season have higher hatch rates and produce male hatchlings which are a rare occurrence during the high rainy season. Although in 2015 the number of nests laid during low season represented just 19.5% of overall nesting, these nests are of high conservation value as they produce the rarer sex and could help population viability. It is important to note that patrols between February and May 2015 were limited due to mechanical problems with the projects quad bike on which patrols are made of the 11 km beach. This resulted in shorter foot patrols during dry season and therefore nesting levels may have been higher than those reported here. Despite this, our study highlights the fact that viable nests are laid year-round and that these nests produce valuable male hatchlings. The majority of these nests are left on the beach and are predated by raccoons and coatis during the first night after laying.

## CONCLUSION

Conservation projects that concentrate effort solely on peak sea turtle nesting season may be inadvertently favouring the production of female hatchlings and leaving male producing nests without protection from illegal take by humans as well as predation by animals. Although it may be tempting to concentrate limited funds to peak season, winter nests are of high value in areas such as Majahuas beach where summer nests do not produce male offspring and are subject to erosion due to tropical storms and hurricanes.

## Authors’ contributions

CEH conceived the study, participated in data collection and analysis and drafted the manuscript; CPLQ carried out the statistical analyses and participated in writing the manuscript; LATS and CEH collected field data; MG analysed the sex ratio data. All authors gave final approval for publication.

## Acknowledgements

We are grateful for the support provided by personnel from the fishers’ cooperative Boca Negra (Cooperativa Pesquera Roca Negra) during our time working on Majahuas beach. Jasiel Noé Juárez-Rábago.

