## Supplemental files for "Neglecting low season nest protection exacerbates female biased sea turtle hatchling production through the loss of male producing nests"

**Supplementary Tables and Figures**

**Temperature effects on sea turtle hatchlings at Majahuas, Mexico**

**Table S1.** Data summary from of nests where female size was obtained.

|  | **Female size** | |  | **Nest data** | | | | |
| --- | --- | --- | --- | --- | --- | --- | --- | --- |
| **Nest ID** | **CCL** | **CCW** | **Season** | **Incubation Date** | **Nest depth** | **Clutch size** | **Hatching success (%)** | **Incubation duration** |
| N4 | 67 | 73 | Dry | 16/02/15 | 39 | 104 | 1.00 | 62.0 |
| N9 | 67 | 73 | Dry | 16/02/15 | 49 | 77 | 0.12 | 62.6 |
| N10 | 69 | 75 | Dry | 16/02/15 | 44 | 92 | 0.92 | 59.9 |
| N11 | 74 | 80 | Dry | 19/02/15 | 45 | 157 | 0.90 | 57.6 |
| N12 | 64 | 69 | Dry | 25/02/15 | 45 | 84 | 0.66 | 62.5 |
| N18 | 67 | 71 | Dry | 25/02/15 | 49 | 97 | 0.92 | 60.7 |
| N21 | 64 | 71 | Dry | 04/03/15 | 45 | 92 | 0.90 | 63.5 |
| N22 | 65 | 76 | Dry | 04/03/15 | 43 | 112 | 0.97 | 56.5 |
| N27 | 64 | 71 | Dry | 19/03/15 | 43 | 89 | 0.85 | 52.9 |
| N28 | 71 | 77 | Dry | 22/03/15 | 51 | 107 | 0.93 | 51.0 |
| N29 | 68.5 | 77 | Rainy | 02/07/15 | ND | 93 | 0.68 | 46.1 |
| N32 | 63 | 68 | Rainy | 05/07/15 | 54 | 106 |  | 49.7 |
| N38 | 71 | 74 | Rainy | 24/08/15 | 57 | 117 |  | 47.9 |

**Supplementary Table 2.** Hatchling morphometry and locomotor test results for *Lepidochelys olivacea* nests at Majahuas beach.

| **Nest ID** | **Season** | **SCL (mm)** | **SCW (mm)** | **Depth (mm)** | **Weight (g)** | **Mean righting propensity** | **Mean of means Righting (s)** | **Run time (min)** |
| --- | --- | --- | --- | --- | --- | --- | --- | --- |
| N1 | Dry | 42.4±0.9 | 36.0±0.8 | 18.3±0.7 | 18.2±0.5 | 5.4 | 3.08 | 2.63 |
| N2 | Dry | 46.44±0.6 | 33.1±0.5 | 24.4±4.1 | 16.6±1.5 | 4.5 | 3.75 | 2.87 |
| N3 | Dry | 39.7±0.8 | 32.6±0.8 | 17.0±1.1 | 15.8±0.8 | 4.5 | 7.47 | 4.61 |
| N4 | Dry | 40.35±0.5 | 31.8±1.8 | 16.1±0.7 | 15.7±1.0 | 3.3 | 6.93 | 2.37 |
| N5 | Dry | 41.1±1.0 | 33.8±1.2 | 18.7±1.3 | 17.2±0.8 | 2.5 | 4.93 | 4.57 |
| N6 | Dry | 39.5±2.3 | 31.7±1.6 | 17.5±1.6 | 15.3±1.5 | 4.2 | 5.41 | 4.01 |
| N7 | Dry | 40.75±0.9 | 33.8±0.7 | 16.9±0.9 | 15.9±0.5 | 5.1 | 3.50 | 2.04 |
| N8 | Dry | 40.7±1.3 | 32.3±0.9 | 17.1±1.2 | 16.7±0.7 | 4.9 | 4.40 | 1.76 |
| N9 | Dry | 40.3±1.1 | 31.5±1.5 | 17.1±1.2 | 15.4±0.7 | 1.0 | 6.53 | 2.31 |
| N10 | Dry | 40.9±0.7 | 32.6±1.3 | 17.2±1.8 | 15.6±0.6 | 5.3 | 2.84 | 1.72 |
| N11 | Dry | 42.3±1.1 | 33.5±1.1 | 20.8±1.0 | 20.8±1.2 | 2.0 | 5.32 | 2.00 |
| N12 | Dry | 39.4±1.4 | 32.4±1.9 | 17.5±0.9 | 15.8±1.6 | 5.1 | 3.56 | 1.23 |
| N13 | Dry | 40.4±0.9 | 34.4±1.5 | 18.5±0.6 | 17.6±0.9 | 5.4 | 3.05 | 1.65 |
| N14 | Dry | 40.5±0.7 | 33.2±0.9 | 17.3±0.8 | 16.6±0.8 | 5.5 | 2.92 | 2.08 |
| N15 | Dry | 40.2±0.8 | 32.5±1.1 | 17.6±0.8 | 15.6±0.6 | 5.0 | 2.64 | 8.77 |
| N16 | Dry | 41.3±0.9 | 33.7±1.1 | 18.6±1.0 | 18.1±0.7 | 0.9 | 5.66 | 4.69 |
| N17 | Dry | 39.9±1.9 | 33.2±1.6 | 16.7±1.0 | 15.8±1.4 | 5.9 | 2.30 | 2.67 |
| N18 | Dry | 40.8±1.1 | 32.2±0.8 | 15.1±0.7 | 14.7±0.6 | 5.6 | 2.50 | 6.54 |
| N19 | Dry | 40.3±1.4 | 34.2±1.2 | 18.0±1.1 | 16.3±1.0 | 5.5 | 3.13 | 1.36 |
| N20 | Dry | 40.1±0.5 | 32.3±0.9 | 17.9±1.0 | 17.4±0.8 | 3.6 | 3.74 | 2.25 |
| N21 | Dry | 40.1±0.9 | 32.6±1.5 | 17.0±0.3 | 15.2±0.7 | 5.1 | 4.15 | 3.28 |
| N22 | Dry | 40.4±1.2 | 31.7±1.0 | 17.2±1.0 | 14.5±0.6 | 4.7 | 3.36 | 1.58 |
| N23 | Dry | 38.8±0.6 | 31.5±1.2 | 17.2±0.8 | 14.9±0.9 | 5.2 | 2.55 | 1.54 |
| N24 | Dry | 38.9±0.8 | 31.2±1.8 | 16.7±0.7 | 13.4±0.8 | 4.4 | 3.38 | 0.95 |
| N25 | Dry | 40.1±0.9 | 34.0±0.9 | 18.4±0.8 | 16.4±0.9 | 5.6 | 3.50 | 1.53 |
| N26 | Dry | 40.5±1.3 | 33.4±0.7 | 17.0±0.7 | 15.3±0.4 | 4.7 | 3.95 | 1.38 |
| N27 | Dry | 40.3±0.8 | 32.7±1.5 | 17.1±0.9 | 15.8±0.8 | 4.7 | 4.09 | 4.28 |
| N28 | Dry | 40.6±2.6 | 30.8±2.5 | 17.0±2.4 | 17.0±0.3 | 3.3 | 4.88 | 2.42 |
| N29 | Rainy | 42.2±0.7 | 33.0±0.8 | 17.2±0.7 | 16.3±1.1 | 5.5 | 3.39 | 3.78 |
| N30 | Rainy | 40.0±4.8 | 33.6±1.1 | 17.3±0.7 | 17.0±0.6 | 5.7 | 2.57 |  |
| N31 | Rainy | 36.3±2.3 | 31.1±2.8 | 15.5±0.9 | 11.5±0.9 | 5.3 | 3.09 |  |
| N32 | Rainy | 39.0±2.0 | 31.8±2.9 | 14.4±0.8 | 11.2±1.4 | ND | ND | ND |
| N33 | Rainy | 42.5±2.2 | 33.2±1.6 | 17.7±1.3 | 17.1±1.6 | 4.1 | 3.61 | 5.17 |
| N34 | Rainy | 41.1±1.0 | 33.0±1.2 | 17.9±0.8 | 15.3±0.5 | ND | ND | ND |
| N35 | Rainy | 37.0±1.1 | 29.7±1.0 | 14.7±2.2 | 12.8±0.5 | 3.1 | 4.41 | 3.25 |
| N36 | Rainy | 41.8±1.6 | 33.4±0.8 | 17.0±0.9 | 15.9±0.8 | 5.8 | 2.32 | 4.21 |
| N37 | Rainy | 40.9±1.1 | 30.4±1.5 | 18.1±1.1 | 16.8±0.5 | 1.2 | 7.41 | 5.06 |
| N38 | Rainy | 39.9±0.7 | 31.0±1.1 | 15.8±1.2 | 15.1±1.6 | ND | ND | ND |

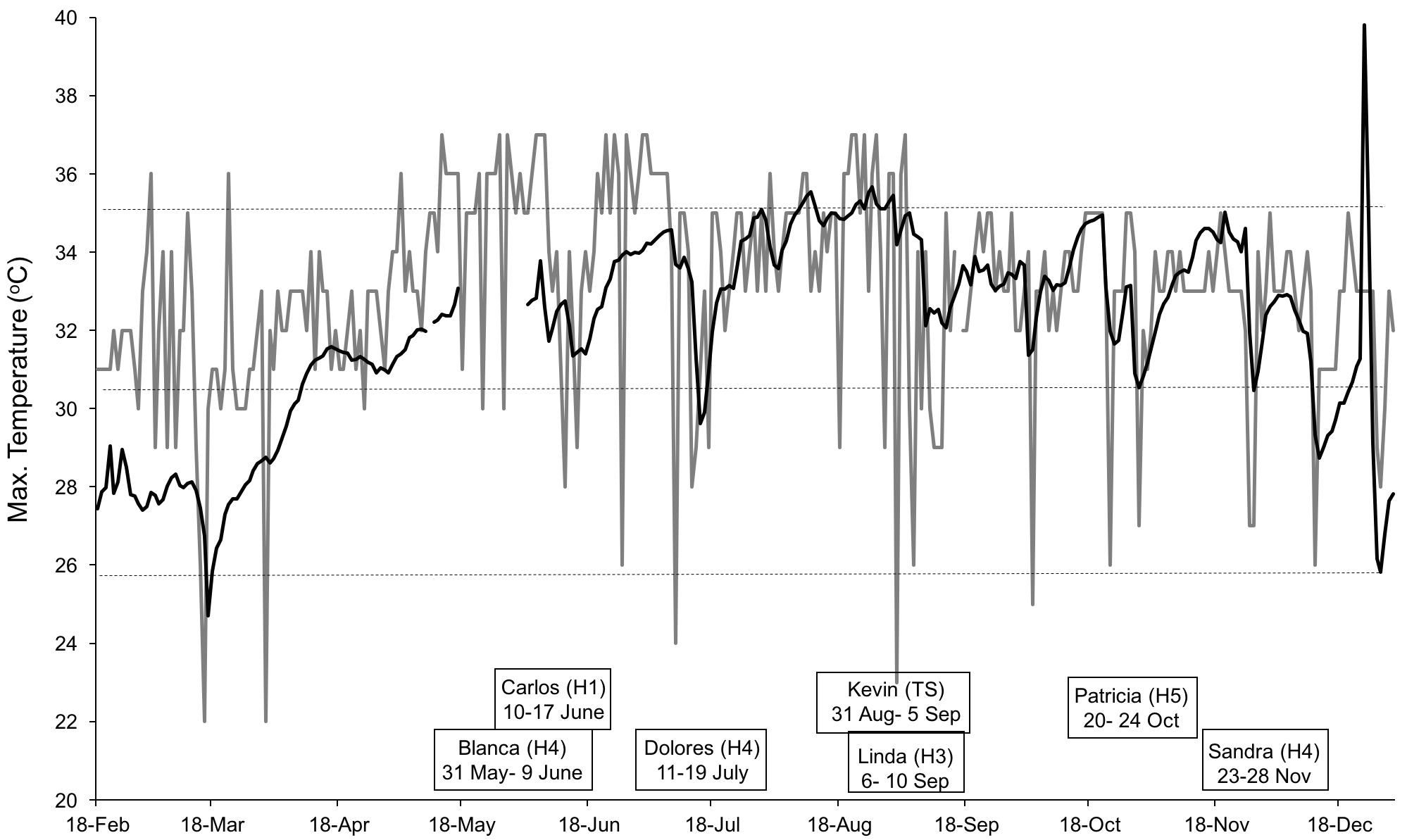

**Supplementary Figure 1.** Maximum air (grey line) and hatchery sand temperature (^o^C) at 45cm depth (black line) between 18^th^ February and 31^st^ December 2015. Boxes contain names of tropical storms (TS) and hurricanes (H + category 1 to 5) that affected the area during this study. Dotted horizontal lines represent the pivotal (30^o^C), and upper (36 ^o^C) and lower viable temperature (26^o^C) for *Lepidochelys olivacea* for the Mexican Pacific region.
